# The small non-coding vault RNA1-1 acts as a riboregulator of autophagy

**DOI:** 10.1101/177949

**Authors:** Rastislav Horos, Anne-Marie Alleaume, Roos Kleinendorst, Abul K. Tarafder, Thomas Schwarzl, Elisabeth M. Zielonka, Asli Adak, Alfredo Castello, Wolfgang Huber, Carsten Sachse, Matthias W. Hentze

## Abstract

Vault RNAs (vtRNA) are small, 88-100nt non-coding RNAs found in many eukaryotes. Although they have been linked to drug resistance, apoptosis and nuclear transport, their function remains unclear. Here we show that a human vtRNA, RNA1-1, specifically binds to the autophagy receptor sequestosome-1/p62. Antisense-mediated depletion of vault RNA1-1 augments, whereas increased vault RNA1-1 expression restricts, autophagic flux in a p62-dependent manner. Bulk autophagy induced by starvation reduces the levels of vault RNA1-1 and the fraction of RNA-bound p62. These findings show that RNAs can act as riboregulators of biological processes by interacting with proteins, and assign a function to a vault RNA.

## Main Text

Eukaryotic small non-coding RNAs, such as miRNAs, siRNAs, tRNAs, and snoRNAs, etc., function as scaffolds that recruit protein complexes to complementary RNA sequences. Whether small non-coding RNAs might have additional functions is less clear.

Vault RNAs (vtRNA) are small non-coding RNA components of giant ribonucleoprotein particles termed vaults (1). Humans express four vtRNA paralogs (vtRNA1-1, vtRNA1-2, vtRNA1-3, vtRNA2-1), which are 88-100nt long and transcribed by RNA polymerase III. Vaults are found in a broad spectrum of eukaryotes ranging from slime moulds to mammals (2). Although vaults can occur at 10,000 to 100,000 particles per cell and have been linked to cellular processes like drug resistance, apoptosis and nuclear transport (3), the function of vaults or vault RNAs remains unclear. Sedimentation experiments showed that only a minor fraction of vtRNAs is incorporated into vaults (4, 5), suggesting they may have functions outside of vault complexes.

Autophagy is an essential process responsible for the recognition, removal and degradation of intracellular components, organelles and foreign bodies within membrane vesicles termed autophagosomes (6, 7). Specific receptors including sequestosome-1/p62 bind ubiquitinated cargos and deliver them to autophagosomes by interaction with LC3 proteins (8). Subsequently, the autophagosomes enclose and fuse with lysosomes to degrade their contents.

Here we show that the autophagy receptor p62 is an RNA-binding protein that binds vault RNAs. We further demonstrate that vault RNA1-1 can control the autophagic flux via p62 at an early stage in the autophagic pathway. Thus, we have identified vault RNA1-1 as a riboregulator of autophagy.

We recently developed a mass spectrometry-based method for the proteome-wide identification of RNA-interacting peptides in RNA-binding proteins (RBPs), termed RBDmap (9). We performed RBDmap on human hepatocellular carcinoma HuH-7 cells and isolated peptides from both known and previously unknown RBPs (10) (Table S1, **Supplementary Materials and Methods**). Among these, we identified a peptide mapping to the zinc finger domain of p62, suggesting that p62 interacts with RNA through this domain. None of the known autophagy receptors have been shown to directly bind RNA (11, 12), and we therefore explored this further.

We verified the p62-RNA interaction by multiple approaches. First, we exposed HuH-7 cells to UV-C light to covalently stabilize direct RNA-protein interactions, and purified crosslinked RNA-binding proteins from lysates using oligo-(dT) coupled beads (13). We confirmed the presence of p62 in the isolated RNP complexes by Western blotting (Fig. 1A). In a complementary approach, we immunoprecipitated (IP) p62 from UV-treated cells after lysis, followed by radioactive labeling of RNA 5’ ends (14). We observed a radioactive signal with a molecular weight corresponding to p62 (Fig. 1B). RNase treatment of the lysates prior to the IP reduced the heterogeneity in band migration, and confirmed the p62-crosslinked entity as RNA (Fig. 1B). Proteomics analysis of the IPs indicated that p62 is the major purified protein at the size of the radioactive signal appearance (Table S2). Therefore, p62 is an RNA-binding protein.

**Fig. 1.**
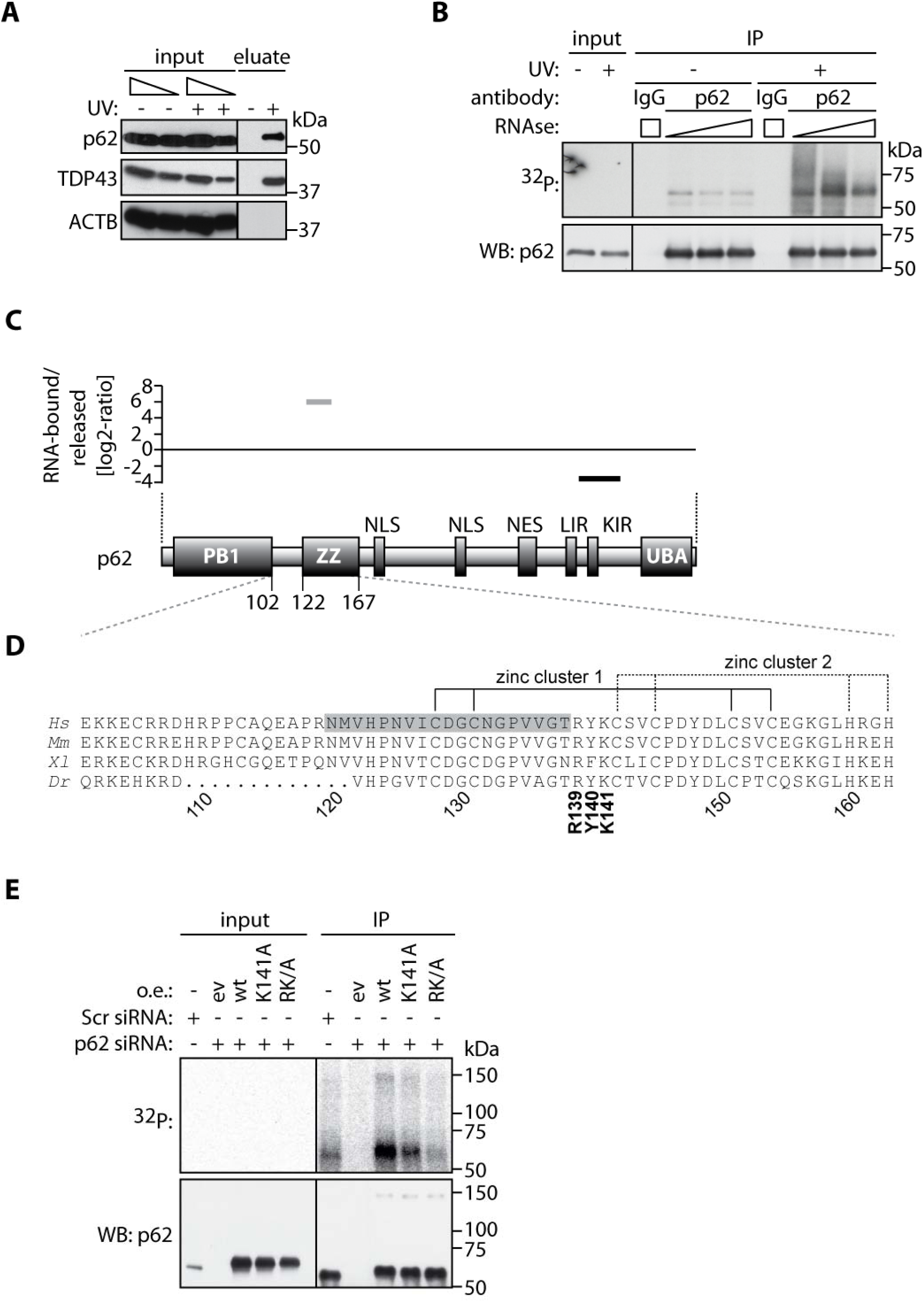
The autophagy receptor sequestosome-1/p62 is an RNA-binding protein. **(A)** Western blot analysis of input and eluate samples from interactome capture experiment. TDP43 serves as a positive control for RNA binding, whereas actin serves as negative control. (**B)** Lysates from UV treated or control cells were treated with dilutions of RNaseA and used for immunoprecipitation followed by radioactive labeling and Western blotting. (**C**) RBDmap-enriched peptide (grey) and a peptide not enriched in the RNA-bound fraction (black) positioned on the p62 protein. The X-axis is scaled to protein length. A scheme of the p62 domain architecture is drawn below. NLS, nuclear localization signal; NES, nuclear export signal, LIR, LC3 interaction region; KIR, Keap1 interaction region; UBA, ubiquitin associated domain. (**D**) Human p62 protein region between AA 101-163. Orthologous proteins are aligned below; dotted region represents insertion of longer peptide. The RBDmap-enriched peptide (FDR 1%) is shaded in grey. *Hs*, *Homo sapiens*, *Mm*, *Mus musculus*, *Xl*, *Xenopus laevis*, *Dr*, *Dario rerio*. (**E**) HuH-7 cells treated with indicated siRNA were transfected with empty vector (ev) or p62 wt and variants (K141A, RK/A refers to the R139/K141-AA). Cells were exposed to 254nm UV-C light, lysed and used for IP followed by the radioactive labeling of RNAs and Western blotting.

To further investigate the p62-RNA interaction, we mutated positively charged or aromatic amino acids within the RNA-binding region implicated by RBDmap (Fig 1C and D). We evaluated tagged p62 variants in HuH-7 cells by IP and RNA labeling. We found that substitution of the conserved residue K141 within the zinc finger domain of p62 appeared to decrease RNA binding (fig. S1A), however the oligomerization of endogenous wild-type p62 with this p62 variant interfered with the analysis. We depleted endogenous p62 from HuH-7 cells by RNAi and, consistent with the above data, the p62-K141A variant showed a substantial reduction in RNA binding compared to p62-wt. RNA binding was further reduced in the R139/K141-AA variant (RK/A, Fig. 1E). These data corroborate the RBDmap result and suggest that the zinc finger domain of p62 is important for its interaction with RNA *in vivo*.

To identify the RNA targets bound by p62 we performed iCLIP, which identifies RNA-protein contacts through crosslink sites (CS) (15). We sequenced RNAs that co-immuno-purified with p62 using two independent antibodies (and the respective controls, fig. S1B and C). All four vault RNAs (vtRNAs) were among the high confidence p62 RNA targets, scoring at the top in the CS density analysis (Table S3) as well as in the RNA class enrichment analysis, respectively (fig. S1D). Differential CS occurrence of individual RNAs isolated from p62 or control IPs, respectively, placed the vtRNAs prominently in the p62 target list (Fig. 2A, **Table S4**). Knock down (KD) of p62 did not affect vtRNAs levels (fig. S1E and F), suggesting that p62 does not regulate vtRNAs stability. Thus, p62 binds a restricted set of RNAs with vtRNAs being the top targets, but it does not mediate vtRNAs degradation.

**Fig. 2.**
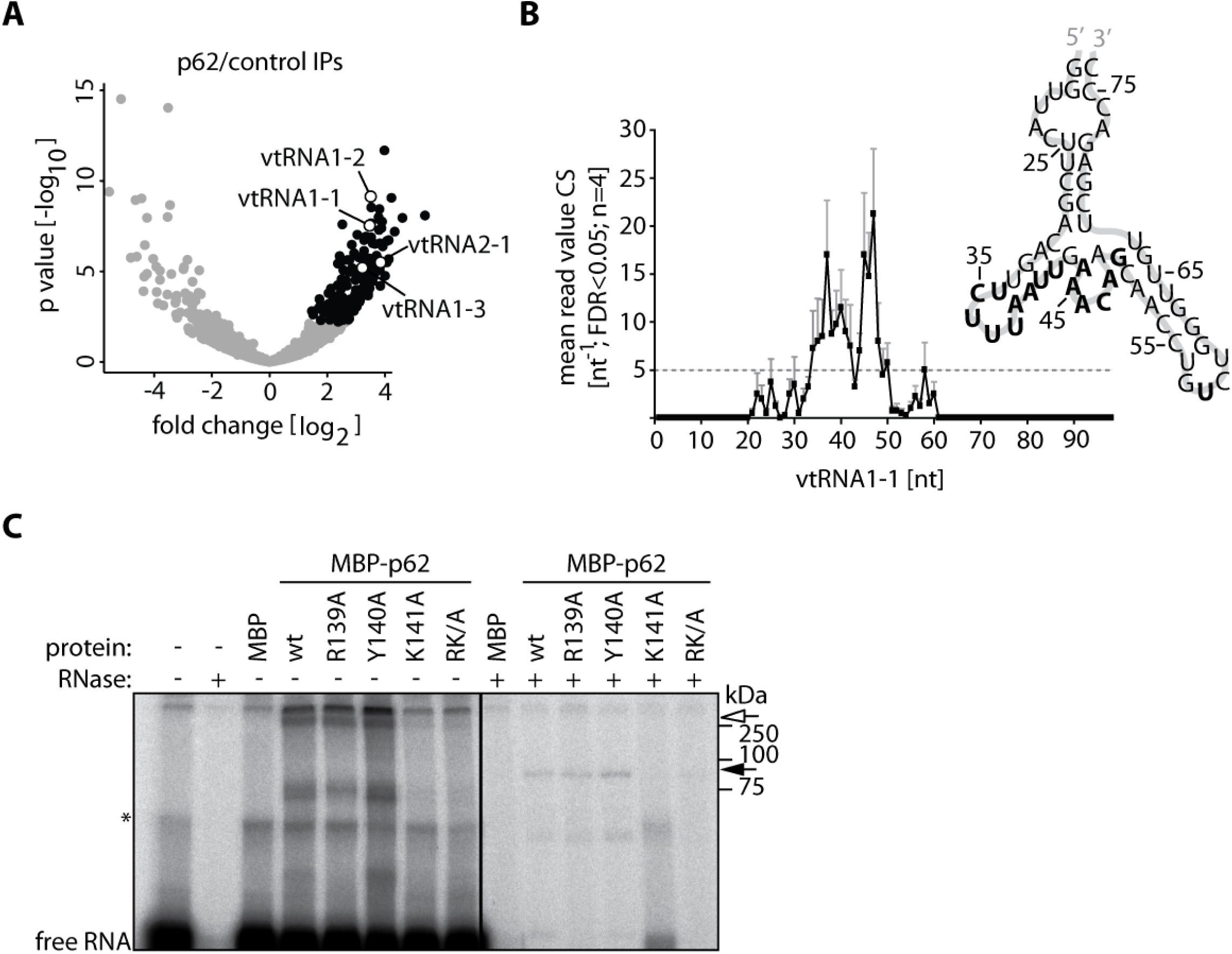
p62 binds vault RNA1-1. (**A**) Volcano plot of genomic regions with differential CS occurrences. The data were normalized for background, and CS enrichment in p62 IPs over controls was tested with DEseq2. The black dots indicate genomic regions (exons, introns) significantly enriched in p62 IPs (p-adj < 0.05). The open circles indicate vault RNAs. (**B**) Significant (FDR 5%) CS read counts of p62 IPs displayed on the vtRNA1-1 transcript sequence. RNA secondary structure is displayed next to the plot and nucleotides with CS mean count values above 5 are indicated in bold. (**C**) Denaturing EMSA using ^32^P-UTP labeled vtRNA1-1 with MBP tag only, MBP-p62 wt or MBP-p62 variant proteins (RK/A refers to the double R139/K141-AA mutant). RNase treatment after the reaction is indicated. Open arrow indicates vtRNA1-1-p62 complex, while the filled arrow indicates RNase-protected fragments at single MBP-p62 unit. * indicates a non-specific band.

Closer inspection revealed that p62 preferentially interacts with looped regions within the central domains of vtRNAs (Fig. 2B and fig. S2). Overexpression of the vtRNA1-1 central domain has previously been shown to confer anti-apoptotic effects independently of vaults (16), therefore we further investigated the interaction of p62 with vtRNA1-1. We established a UV crosslinking and electrophoretic mobility shift assay (EMSA) to evaluate the p62-vtRNA1-1 interaction *in vitro*. Incubation of radiolabeled vtRNA1-1 and MBP-tagged p62 led to the formation of labeled, higher molecular weight RNP complexes (Fig. 2C). We found that the K141A and RK/A substitutions reduced the formation of these RNP complexes, consistent with the *in vivo* observations. RNase treatment after crosslinking diminished non-specific interactions, resulting in a protected RNA fragment bound to monomeric MBP-p62 (90 kDa) (Fig. 2C). The K141A and RK/A substitutions abolished this RNase-resistant RNA fragment. These findings mirror the analysis of p62 mutants *in vivo*, and suggest that the p62-vtRNA1-1 interaction requires residues within p62 zinc finger domain.

To explore a possible function of the p62-vtRNA1-1 interaction in autophagy, we overexpressed vtRNA1-1 in HuH-7 cells and monitored autophagic flux by assessing LC3B conjugation from LC3B I to LC3B II during autophagosome assembly and p62 levels, reflecting autolysosomal degradation. Increasing vtRNA1-1 levels reduced LC3B conjugation and induced p62 protein accumulation in a dose-dependent manner (Fig. 3A and S3A), suggesting decreased autophagic flux. Overexpression of other vault RNAs did not significantly affect the LC3B-II/LC3B-I ratio (Fig. 3B and S3A). Treatment with bafilomycine A_1_ (BafA), an inhibitor of autophagosome-lysosome fusion that leads to the accumulation of mature autophagosomes, restored the LC3B conjugation ratio in cells overexpressing vtRNA1-1 (Fig. 3B). This result suggests that vtRNA1-1 overexpression did not disturb autophagosome turnover but rather restricted autophagic flux. Further, LNA-mediated KD of vtRNA1-1 resulted in a dose-dependent decrease in p62 levels and increased LC3B conjugation (Fig. 3C and S3B). Concurrent removal of p62 (Fig. 3D, compare lanes 3 and 4 with 1 and 2) or BafA treatment (Fig. 3D, compare lanes 5 and 6 with 1 and 2) restored the LC3B conjugation ratio in vtRNA1-1 KD cells as compared to control cells. These experiments suggest that the removal of vtRNA1-1 did not inhibit autophagosome turnover and that vtRNA1-1 affects early stages of autophagy via p62.

**Fig. 3.**
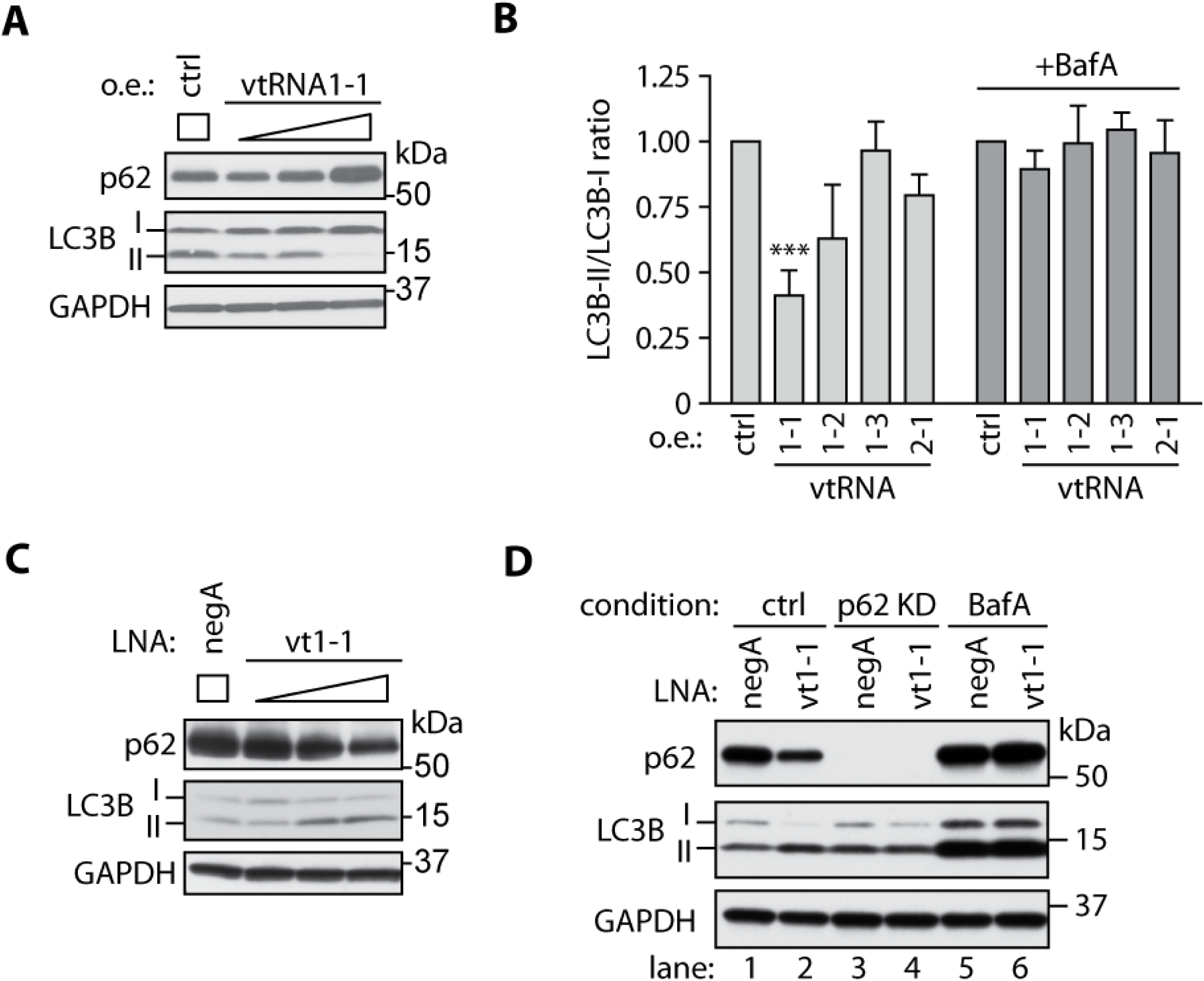
vtRNA1-1 regulates autophagy via p62. (**A**) HuH-7 cells were transfected with empty vector (ctrl) or increasing amount of vector encoding vtRNA1-1 and lysed after 24 hours. Lysates were analyzed by Western blot with the indicated antibodies. (**B**) Cells were transfected with indicated vtRNAs, vehicle treated or treated with BafA at 100 nM for 5 hours, and then lysed. Lysates were analyzed by Western blotting and images of LC3B staining were quantified. n=3, *** *p*<0.005 (**C**) Cells were transfected with a control LNA oligo (negA), or with increasing amounts of LNA oligos targeting vtRNA1-1, and lysed after 48 hours. Lysates were analyzed by Western blot with the indicated antibodies. (**D**) Cells were transfected with indicated LNA oligos and control or p62 siRNA and incubated for 48 hours. Where indicated, cells were treated with BafA. Lysates were analyzed by Western blotting with the indicated antibodies.

Next, we investigated the role of the p62-vtRNA1-1 interaction during bulk cytosol autophagy induced by amino acid and serum starvation. During bulk autophagy, p62 supports the increased autophagic flux (17), and also serves as degradation substrate (18). Indeed, cells that are starved in the presence of BafA show pronounced expression and co-localization of autophagosomal LC3B and p62 (fig. S4A). We found that the p62-RNA interaction gradually decreases during 6 hours of starvation and that this reduction is exacerbated by BafA treatment (Fig. 4A and B). These data indicate that bulk autophagy decreases the fraction of RNA-bound p62 relative to total p62, and that p62 destined for lysosomal degradation no longer binds RNA. Consistent with this hypothesis, we found that RNA-bound p62 is enriched in the detergent-soluble subcellular fraction and depleted from the vesicular/nuclear and membrane debris fractions (Fig. 4C). For p62, these data reveal an anti-correlation between engagement with RNA and engagement with the autophagy apparatus, suggesting that RNA binding interferes with p62’s function during autophagosome formation at an early stage of autophagy.

**Fig. 4.**
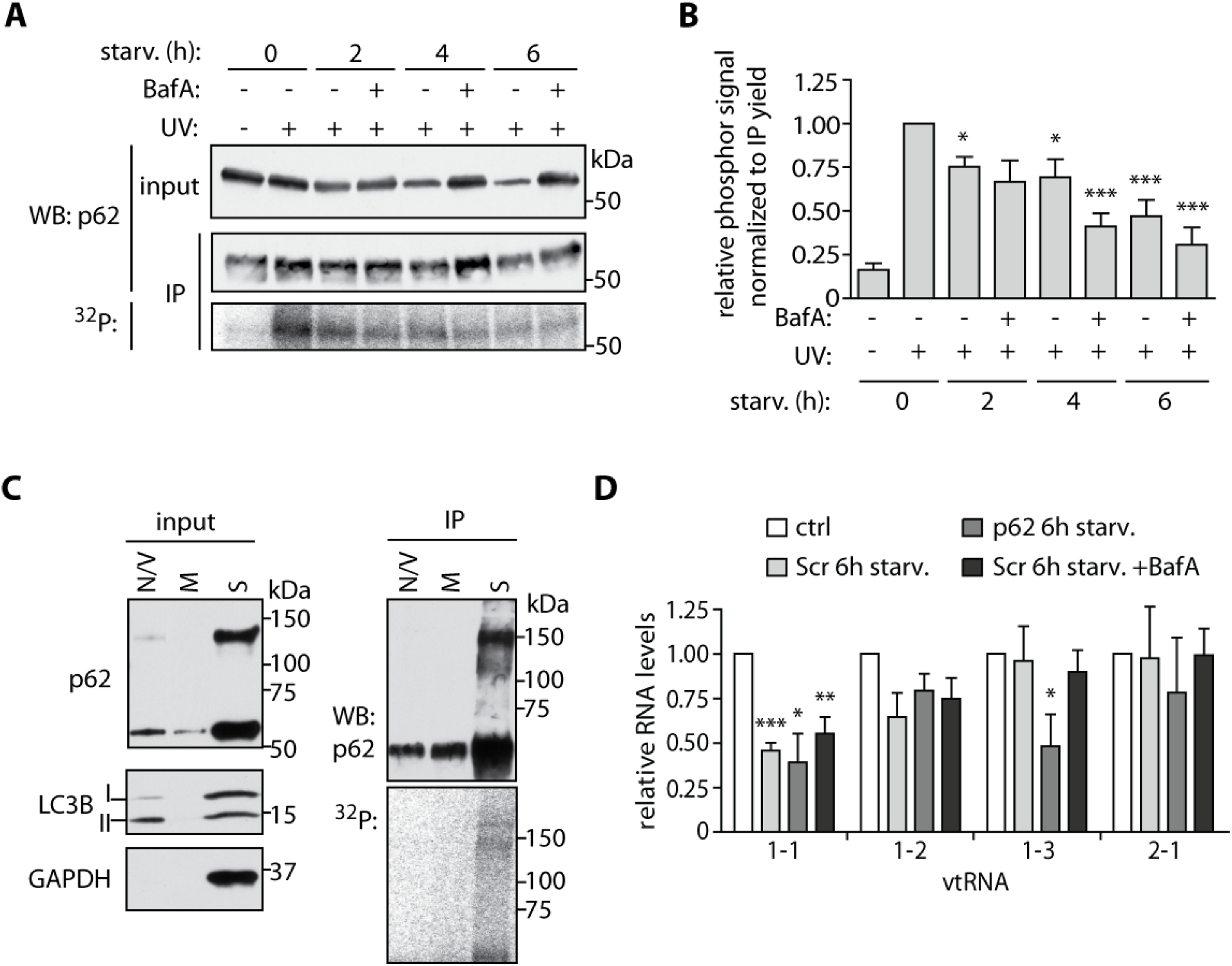
Starvation reduces p62 RNA binding and vtRNA1-1 expression. (**A**) Cells were starved in minimal medium containing solvent control or BafA at 100 nM for the indicated time, 254nm UV-C light exposed and lysed. Lysates were used for p62 IP and RNA radiolabeling assay. After SDS-PAGE and transfer, the membrane was exposed overnight on film and used subsequently for Western blotting. (**B**) Phosphorimages and Western blots of 3 independent replicates as described in (A) were used for quantification and ratio calculation (radioactive/IP Western blotting signal). n=3, * *p*<0.05, *** *p*< 0.005 (**C**) HuH-7 cells were exposed to 254 nm UV-C light and lysed in hypotonic buffer. A nuclear and vesicular fraction (N/V) was collected, followed by the pelleting of membranous debris (M). Both pellets and the supernatant (S) were then used for p62 IP and RNA radiolabeling assay. (**D**) HuH-7 cells were treated with a scrambled control siRNA (Scr) or p62 siRNA for 48 hours, followed by starvation for 6 hours. Total RNA was isolated and analyzed by Northern blotting; phosphorimages were quantified and normalized with the tRNA Gln^*CUG*(3-2)^ to the 0 hour time point, n=3, * *p*<0.05, ** *p*<0.01, *** *p*< 0.005

In addition, the levels of vtRNA1-1, but not other vault RNAs, drop significantly in HuH-7 cells after 6 hours of starvation (Fig. 4D). The starvation-induced decrease in vtRNA1-1 levels is not a result of co-degradation with p62, because neither the KD of p62 nor the treatment with BafA significantly restored vtRNA1-1 levels (Fig. 4D). The decrease in vtRNA1-1 levels correlates with and possibly causes a decrease in the fraction of RNA-bound p62 during starvation-induced autophagy (fig. S4B). Of relevance, PolIII transcription at the vtRNA1-1 locus is dynamically controlled by its repressor MAF1, which in turn is activated by starvation (19), suggesting that starvation represses the transcription of vtRNA1-1. Thus, bulk autophagy leads to a concurrent decrease in vtRNA1-1 levels and the fraction of p62 associated with RNA.

Here we show that the small non-coding RNA vtRNA1-1 regulates autophagic flux through the autophagy receptor p62, assigning a function to the first member of this enigmatic family of non-coding RNAs described more than 30 years ago (1). Future work will address the functions of the interaction of the other members of the vault RNA family with p62. Moreover, it will be illuminating to unravel the mechanistic and structural details of how vtRNA1-1 controls the function of p62 in autophagy. Riboregulation of protein function may represent a new general paradigm, complementing well-established forms of regulation such as by protein-protein interactions.

## Acknowledgments

The sequencing data were deposited in Array Express with the accession number E-MTAB-4894. This work was supported by a European Research Council Advanced Grant and by a grant from the Virtual Liver Consortium (German Ministry for Education and Research) (to M.W.H.). We are grateful to Bernd Fischer (German Cancer Research Center, Heidelberg, Germany) for the analysis of the RBDmap dataset, to Bernd Klaus (Centre for Statistical Data Analysis, EMBL) for help with statistical analyses, to Charles Girardot (Genome Biology Computational Support, EMBL) for help with sequencing data deposition, to Mandy Rettel (Proteomics facility, EMBL) for proteomic analyses, and to members of the Hentze group for critical reading of the manuscript. The authors declare no competing financial interests.

## Supplementary Material

### Materials and Methods

#### Cell culture and chemicals

HuH-7 cells were cultured in low glucose (5mM) DMEM supplemented with 10% heat inactivated FCS (PAA), 2mM L-glutamine (25030081, Thermo Fisher) and 100 U/ml PenStrep (15140122, Thermo Fisher). We derived a HuH-7 Flp-In TREx cell line using published protocols (Flp-In T-Rex, Thermo Fisher), and prepared stably expressing doxycycline-inducible cell lines following manufacturer’s instructions. Stable cell lines were grown in medium containing blasticidine (5µg/ml) and zeocin (100 µg/ml) or hygromycin (200 µg/ml). Induction was performed with doxycycline at 100 ng/ml overnight. Transfections were done using Lipofectamine 3000 (L3000008, Thermo Fisher) for the plasmids, or Lipofectamine RNAiMax (13778075, Thermo Fisher) for the siRNA and LNAs. Bafilomycine A_1_ (tlrl-baf1, InvivoGen) was diluted in DMSO to 100 µM and used at 50-100 nM for 4-6 hours. 4-thiouridine (T2933, Biomol) was used at 100 µM for 16 hours. For starvation, cells were washed twice with PBS and starved in low glucose DMEM lacking amino acids (D9800-13, USBiological) and serum.

#### RNA isolation and Northern blotting

RNA was isolated using TRI reagent (T9424, Sigma-Aldrich) as recommended by the manufacturer. RNA was dissolved in nuclease-free water and stored at −80°C. Typically, 10 or 15 µg of total RNA was mixed with 2x loading dye (95% formamide; 0.025% xylene cyanol and bromophenol blue; 18mM EDTA; 0.025% SDS), denatured for 5 min at 95°C, cooled on ice and loaded on 8% acrylamide (19:1), 7M urea polyacrylamide gels. A semi-dry blotting apparatus was used for blotting on Hybond N^+^ membranes (RPN1520B, GE) which were UV auto-crosslinked, pre-hybridized for 1 hour at 50°C and used for hybridizations with ^32^P labelled DNA antisense oligonucleotide probes overnight at 50°C. The membranes were then washed three times with high stringency buffer (5X SSC; 5% SDS), three times with low stringency buffer (1X SSC; 1% SDS) and exposed to phosphorimaging screens for 4 hours or overnight. Screens were scanned at Typhoon FLA-7000 (GE) and TIFF images were quantifiied by ImageJ.

#### RNA interactome capture

RNA interactome capture was performed with minor modifications in the cell lysis procedure as previously described (S*1*). The cells were washed twice with PBS on ice before UV crosslinking at 150 mJ/cm^2^. Cells were lysed directly with lysis buffer on the cell culture plates, scraped and lysates were sheared through a 27G needle before incubation with oligo d(T) beads (volume ratio lysate to beads 15:1) for 1 hour at 4°C. Beads were then washed twice with each wash buffer, and pooled eluates from three rounds of purification were used for RNAse treatment, concentration using Amicon 3K columns (UFC500396, Merck Millipore) and mixing with 4x sample buffer (4xSB) (200mM Tris-HCl pH6.8; 8% SDS; 40% Glycerol, 0.04% bromophenol blue, 400mM DTT; 10% beta mercaptoethanol) for SDS-PAGE.

#### RBDmap

RBDmap for HuH-7 cells was performed and analyzed as described (S*2*). The data can be accessed at http://www.hentze.embl.de/public/RBDmapHuH7.

#### Protein extracts, SDS-PAGE and Western blotting

For Western blotting, cells were washed twice with ice cold PBS on ice and lysed on plate using RIPA lysis buffer (89900, Thermo Fisher) supplemented with protease inhibitor (11873580001, Roche). Lysates were treated with benzonase (100U/ml, 71206, Merck Millipore) for 15 min on ice and the protein concentrations were measured. Lysates were mixed with 4xSB, boiled for 5 min and typically 15 µg of lysate was used for SDS-PAGE. Proteins were transferred to nitrocellulose or PVDF membranes using the Trans-Blot Turbo Transfer System (Bio-Rad) and blocked for 1 hour at room temperature with 5% milk in PBS; 0,05% Tween (PBS-T). Primary antibodies were incubated in 5% milk PBS-T either overnight at 4°C or 1 hour at RT, followed by 3x PBS-T washes, secondary antibody incubation in 5% milk in PBS-T for 1 hour at RT, 3x PBS-T washes and developed using ECL (WBKLS0500, Millipore). Antibodies used were anti-p62 (1:20 000; PM045, MBL; 1:20 000; H00008878-M01, Novus), TDP43 (1:10 000; 10782-2-AP, ProteinTech Group), β-actin (1:20 000; A5441, Sigma-Aldrich), LC3B (1:20 000; PM036, MBL) and GAPDH (1:20 000; G9545, Sigma-Aldrich). Secondary antibodies (goat anti-mouse IgG-HRP, sc-2005; goat anti-mouse IgG-HRP, sc-2004, Santa Cruz) were used at 1:20 000 dilution.

#### siRNA, LNAs

An siRNA pool targeting p62 (L-010230-00-0020, GE) was used at 30 nM concentration for 48 hours. As control siRNA an equimolar mix of Scramble (5’ UUCUCCGAACGUGUCACGUtt 3’; s229174, Thermo Fisher), sLuciferase (5’ CGGAUUACCAGGGAUUUCAtt 3’; Thermo Fisher) and SWNeg9 (5’ UACGACCGGUCUAUCGUAGtt 3’; s444246, Thermo Fisher) was used. LNAs (Exiqon) targeting vtRNA1-1 (#1: 5’ ttaaagaactgtcgaa 3’; #3: 5’ttaaagaactgtcga 3’) and control negA (5’ aacacgtctatacgc 3’) were used at 25 or 50 nM for 48 hours.

#### Immunoprecipitations (IP)

0.75 µg of p62 antibody or appropriate control IgG was coupled for 1 hour at RT to 12.5 µl of Protein G coupled magnetic beads (10004D, Thermo Fisher). Cells were washed twice with cold PBS, lysed in lysis buffer (100mM NaCl; 50mM Tris-HCl pH7.5; 0.1% SDS; 1 mM MgCl_2_; 0.1 mM CaCl_2_; 1% NP40; 0.5% sodium deoxycholate; protease inhibitors (11873580001, Roche)) and homogenized by ultrasound (level 4, 3x 10sec, 50% amplitude) on ice. Lysates containing 2 mg of total protein were used for IP for 1 hour at 4°C, washed three times with high salt buffer (500mM NaCl; 20mM HEPES pH7.3; 1% NP-40; 0.1% SDS; 1 mM EDTA; 0.5% sodium deoxycholate; protease inhibitors (11873580001, Roche)) and three times with the lysis buffer. Proteins were eluted at low pH (0.1M glycin pH2.0) and neutralized with 0.2M Tris-HCl pH8.5.

#### Proteomics

**In gel digestion.** For in-gel processing, eluates from p62 and control IPs were separated by SDS-PAGE and stained with coomassie blue. Gel slices from the region around 60 kDa were cut from the gel and subjected to in-gel digestion with trypsin (S*3*). Subsequently, peptides were extracted from the gel pieces. Samples were sonicated for 15 minutes, centrifuged and the supernatant removed and placed in a clean tube. Subsequently, a solution of 50:50 water: acetonitrile, 1 % formic acid (2 × the volume of the gel pieces) was added and the samples were again sonicated for 15 minutes, centrifuged and the supernatant pooled with the first. The pooled supernatants were then dried down with the speed vacuum centrifuge. The samples were dissolved in 10 µL of reconstitution buffer (96:4 water: acetonitrile, 0.1% formic acid and analyzed by LC-MS/MS.

**LC-MS/MS – Dionex LC.** Peptides were separated using the UltiMate 3000 RSLC nano LC system (Dionex) fitted with a trapping (Dionex Acclaim PepMap100, 75 µm × 2 cm, C18, 3 µm, 100 Å) and an analytical column (Dionex Acclaim PepMap RSLC 75 µm × 15 cm C18, 2 µm, 100 Å). The outlet of the analytical column was coupled directly to a Q-Exactive (Thermo) using the proxeon nanoflow source in positive ion mode. Solvent A was water, 0.1 % formic acid and solvent B was acetonitrile, 0.1 % formic acid. The samples were loaded using the µLpickup mode of the autosampler, with a constant flow of solvent A at 6 µL/min onto the trapping column. Trapping time was 5 minutes. Peptides were eluted via the analytical column a constant flow of 0.3 µL/min. During the elution step, the percentage of solvent B increased in a linear fashion from 4 % to 7 % B in 5 minutes, then from 7 % to 25 % in a further 30 minutes and finally from 25 % to 40 % in another 5 minutes. Column cleaning at 85 % B followed, lasting 5 minutes, before returning to initial conditions for the re-equilibration, lasting 10 minutes.

**QE – MS - DDA.** The peptides were introduced into the mass spectrometer (Q-Exactive, Thermo) via a Pico-Tip Emitter 360 µm OD × 20 µm ID; 10 µm tip (New Objective) and a spray voltage of 1.8 kV was applied. The capillary temperature was set at 250 °C. Full scan MS spectra with mass range 300-1500 m/z were acquired in profile mode in the FT with resolution of 70000. The filling time was set at maximum of 32 ms with a limitation of 1x106 ions. DDA was performed with the resolution of the Orbitrap set to 17500, with a fill time of 60 ms and a limitation of 5x105 ions. Normalized collision energy of 25 was used. A loop count of 15 with count 1 was used. Dynamic exclusion time of 30s was applied. An underfill ratio of 1%, corresponding to 8.3 x104 ions was used. The peptide match algorithm was set to ‘off’ and only charge states of 2+, 3+ and 4+ were selected for MS/MS. Isolation window was set to 2 m/z and 110 m/z set as the fixed first mass. MS/MS data was acquired in centroid mode. In order to improve the mass accuracy, a lock mass correction using a background ion (m/z 445.12003) was applied.

#### Polynucleotide kinase (PNK) assays

After homogenization the lysates were treated with 10 ng/µl of RNase A (R5503, Sigma-Aldrich) and 2U/ml Turbo DNAse (AM2238, Thermo Fisher) for 15 min at 37°C, cooled on ice and used for IPs. After the IP and washes, beads were washed additionally with PNK buffer (50mM NaCl; 50mM Tris-HCl pH7.5; 10mM MgCl2; 0.5% NP-40; protease inhibitors (11873580001, Roche)), then resuspended in PNK buffer containing 0.1 µCi/µl [?-32P] rATP (Hartmann), 1 U/µl T4 PNK (NEB), 1mM DTT and labeled for 15 min at 37°C. After 4 washes with PNK buffer, proteins were eluted as described above, resolved by SDS-PAGE, blotted and membranes were exposed overnight to phosphorimager screens or to the imaging film (Z350397-50EA, Sigma), followed by Western blotting.

#### iCLIP

iCLIP was performed as published (S*4*) using IPs described above. Treatment of the lysates with RNAseI (AM2295, Thermo Fisher) was used at 20 U/ml.

#### Bioinformatics and statistical analyses

The analysis of the p62 iCLIP datasets is described at http://www.hentze.embl.de/public/p62-iCLIP. RNA secondary structures were predicted using the ViennaRNA package. Data are displayed as mean ± SEM, and two-tailed unpaired t test was used. Images were quantified with ImageJ.

#### Cloning

Full length human p62 wild type cDNA was cloned into pcDNA5_FRT/TO vector with N-terminal FLAG/HA tag (MDYKDDDDKSAGGYPYDVPDYAKL…) using HindIII and XhoI sites. Single and double amino acid mutations were done using PCR-mediated mutagenesis. Recognition sites of p62 siRNA were mutated in synonymous fashion (5’ GGATCGAGGTAGACATAGA 3’; 5’ GAGCAAATGGAATCCGACA 3’; 5’ GGACGCACCTCTCATCTAA 3’; 5’ CGACTGGCCTCAAAGAGGC 3’), cDNA was synthetized in pUC57 (GenScript) and swapped into p62 cDNA using BamHI and XhoI sites. Vault RNA with T7 or H1(2xTO) promotors were synthetized (GenScript) in pUC57 backbone.

#### In vitro transcription, RNA quantification and EMSA

pUC57 plasmid with T7_vault RNA1-1 was used for in vitro transcription reaction using MEGAshortscript kit (AM1354, Thermo Fisher) with ^32^P-αUTP (SRP-210, Hartmann) according to the manufacturer’s protocol. RNA was gel purified, phenol-chloroform extracted, dissolved in water and measured for the specific activity with scintillation counter and concentration with QuBit (Thermo Fisher). EMSA reactions containing 1 µM of proteins, 30 nM of RNA, 50 ng/µl of BSA and reaction buffer (50mM KCl; 10mM HEPES pH7.3; 0.25mM EDTA; 2,5mM MgCl2; 5%glycerol; 0.1% NP-40; 1mM DTT) were incubated 20 min at room temperature. After the reaction, heparin was added (final concentration 100 ng/µl) and samples were exposed to 10 min (corresponding to 1500 mJ/cm^2^) of 254 nm UV-C light on ice. RNaseA (final concentration 100ng/µl) treatment was performed 15 min at 37°C where indicated. Samples were then mixed with 4xSB, incubated at 70°C for 10 min and analyzed by denaturing SDS-PAGE. Gel was dried for 1 hour at 80°C and exposed overnight to phosphorimager screen.

#### p62 protein expression and purification

MBP-p62-his_6_ was expressed by autoinduction in ZY media for 16 hrs at 20°C. Cells were lysed by resuspension in lysis buffer (50mM HEPES pH 8.0, 1M NaCl, 0.5 mM TCEP, 1x protease inhibitor) followed by four passes through a microfluidizer. Lysate was clarified by centrifugation at 48 000 *g* and incubated with Ni-NTA beads for 1 hr. Beads were washed extensively in buffer 1 (50mM HEPES pH 8.0, 1M NaCl, 0.5 mM TCEP, 50mM Imidazole) and protein eluted with buffer 2 (50mM HEPES pH 8.0, 1M NaCl, 0.5 mM TCEP, 250mM Imidazole).

#### Immunofluorescence microscopy

For immunostaining, cells were cultured on ibidi slides (80426; ibidi), fixed for 10 min with 4 % paraformaldehyde, washed with PBS, permeabilized and blocked for 30 min in 0.1% Triton-X 100 in 1% BSA solution. Cells were then incubated with primary antibodies for 2 hours at room temperature, washed in PBS and incubated with the secondary antibody and DAPI for 1 h at room temperature in the dark. Slides were washed 3 times in PBS and stored at 4°C in PBS until imaging. Reagents used were anti-p62 1:500 (PM045, MBL), anti-LC3B 1:300 (CTB-LC3-2-IC, Cosmo Bio), anti-mouse IgG Alexa Fluor 488 1:1000 (4408, Cell Signaling), anti-Rabbit IgG Alexa Fluor 555 1:1000 (4413, Cell Signaling), DAPI (10236276001 Roche). Fluorescent staining was viewed on a wide-field fluorescence microscope (Cellobserver HS; Carl Zeiss) equipped with a 63x Plan-Apochromat 1.4 oil objective and a Lumen Dznamics 120 LED light source. For detection of Dapi (Ex 353nm / Em 465nm), Alexa Fluor 488 (Ex 493nm / Em 517nm) and Alexa Fluor 555 (Ex 553nm / Em 568nm), the AxioCamMRm3 camera was used. Pictures were acquired using Zeiss software (ZEN 2 blue edition) and exported by ImageJ (version 2.0.0-rc-41/1.50d).

#### Sub-cellular fractionation

Cells were washed twice with PBS on ice before 254nm UV-C light exposure with 150 mJ/cm2. Cells were lysed directly on the plates with hypotonic buffer (10mM HEPES pH7.3; 20mM KCl; 1mM EDTA; 1% triton X-100; 1mM DTT; protease inhibitors (11873580001, Roche)) by swelling 10min on ice, followed by scraping. Lysates were homogenized by gentle pipetting and centrifuged for 10 min at 2.500xg at 4°C. The nuclei/vesicles-containing pellet (P2.5) was carefully washed in hypotonic buffer, resuspended in extraction buffer (20mM HEPES pH7.3; 200mM NaCl; 5mM MgCl_2_; 1% NP-40; protease inhibitors), sonicated, treated with 2U/ml Turbo DNase and RNaseA (see PNK assays procedure), salt content was adjusted by addition of 5xIP buffer (100mM HEPES pH7.3; 1M NaCl; 5mM EDTA; 5% NP-40; 0.5% SDS; 2.5% sodium deoxycholate; protease inhibitors) and samples were processed for IP. The supernatant (S2.5) was further centrifuged for 20 min at 20.000xg at 4°C. The membrane debris containing pellet (P20) was resuspended in the extraction buffer, sonicated, treated with 2U/ml Turbo DNase and RNaseA, salt content was adjusted by addition of 5xIP buffer and samples were processed for IP. The supernatant (S20) was treated with 2U/ml Turbo DNase and RNaseA, salt content was adjusted by addition of 5xIP buffer and samples were processed for immunoprecipitation.

**Figure S1.**
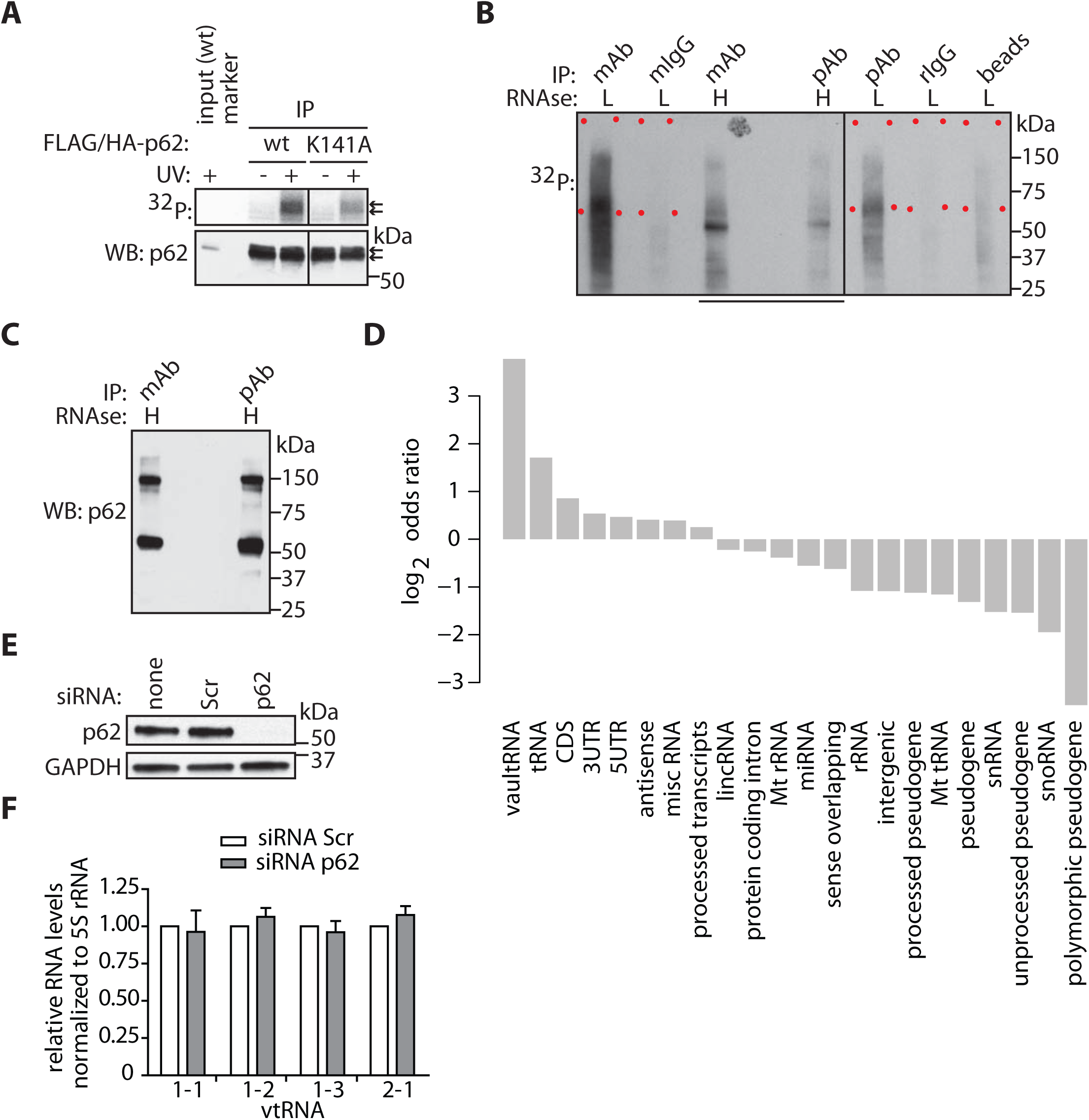
Analysis of p62 RNA binding. (**A**) Stable HuH-7 cell clones expressing inducible FLAG/HA-tagged p62 wt and K141A variant were exposed to 254nm UV-C, lysed and IP with anti-HA antibody-conjugated beads and radioactive labeling was performed. The blot was exposed to phosphorimager screen and subsequently used for Western blotting. Endogenous and exogenous p62 are indicated by arrows. (**B**) Lysates from 254nm UV-C light exposed HuH-7 cells were treated with low (20U/ml RNaseI, L) or high concentration of RNase (200 U/ml, H), and used for IPs with indicated antibodies and controls. p62-RNA complexes were separated by SDS-PAGE, blotted and excised as indicated by the red dots rectangles. The underlined area of the blot was used for a subsequent Western blotting shown in panel (**C**). (**D**) Log_2_ ratios of RNA enrichment in p62 IPs over the control IPs (Fisher exact test, *p-adj* < 0.05). (**E**) Cells were transfected with siRNA for 48 hours, then lysed and analyzed by Western blotting with the indicated antibodies. (**F**) Total RNA was isolated and analyzed by Northern blotting. vtRNA probe signals were quantified and normalized to the 5S rRNA, n=3.

**Figure S2.**
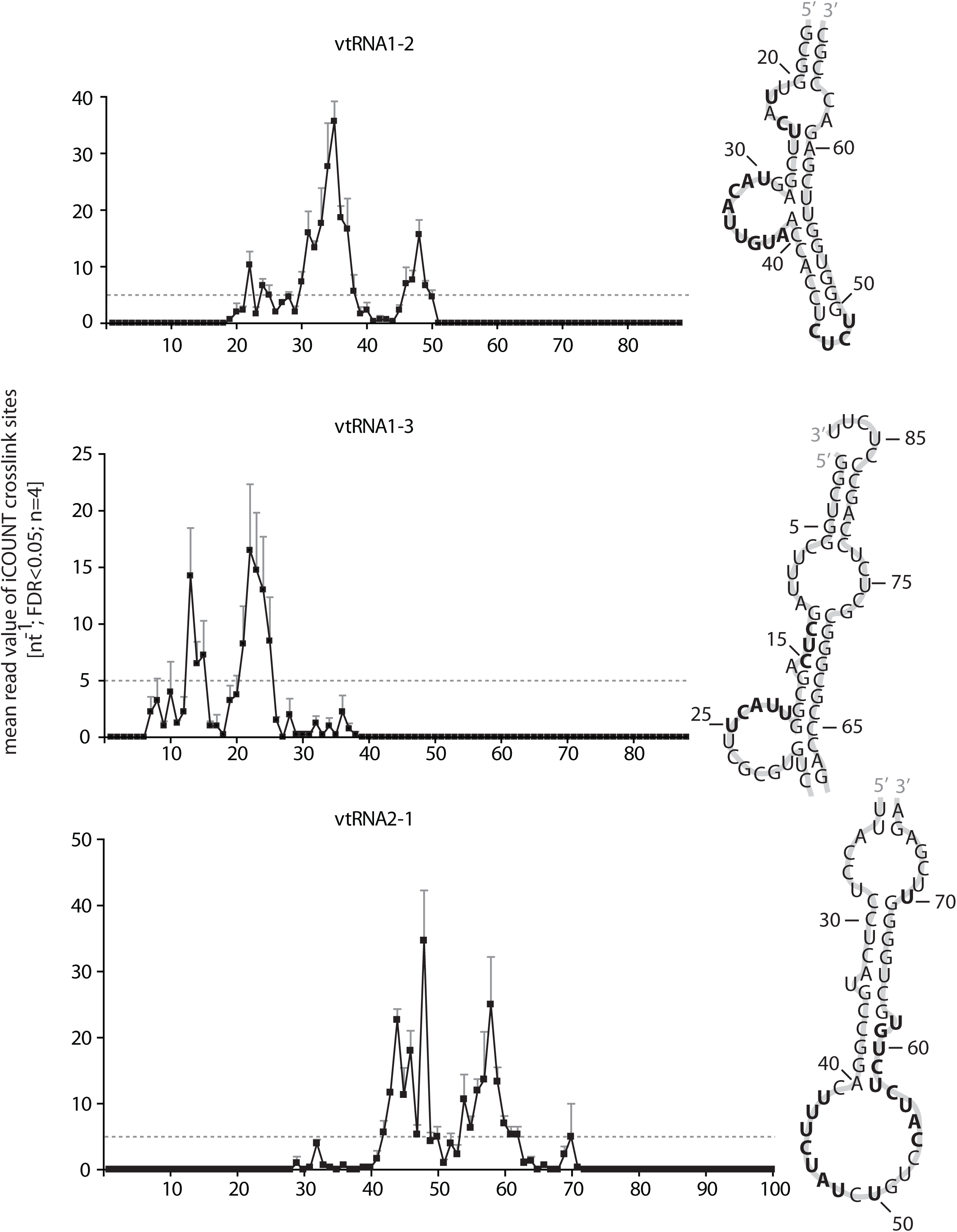
p62 crosslink-sites (CS) analysis on vtRNAs. Significant (FDR 5%) CS read counts of p62 IPs displayed on the vtRNAs transcript sequence. RNA secondary structure is displayed next to the plot and nucleotides with CS mean count values above 5 are indicated in bold.

**Figure S3.**
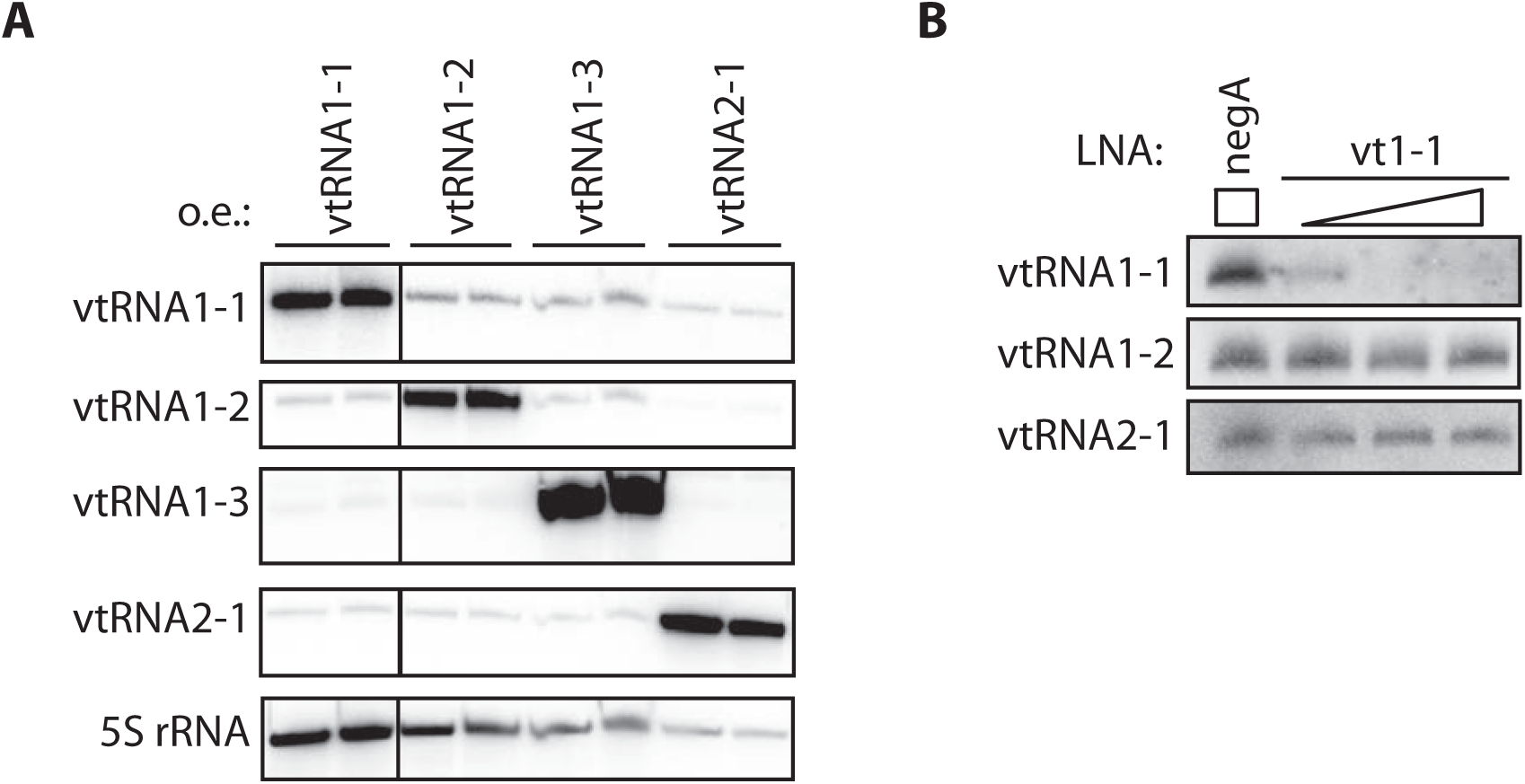
Overexpression and knock-down of vtRNAs. (**A**) Cells were transfected with plasmids expressing vault RNAs and lysed after 24 hours. Total RNA was isolated and analyzed by Northern blotting. (**B**) Cells were transfected with control LNA oligo (negA) or increasing amounts of LNA oligo targeting vtRNA1-1 and lysed after 48 hours. Total RNA was isolated and analyzed by Northern blotting.

**Figure S4.**
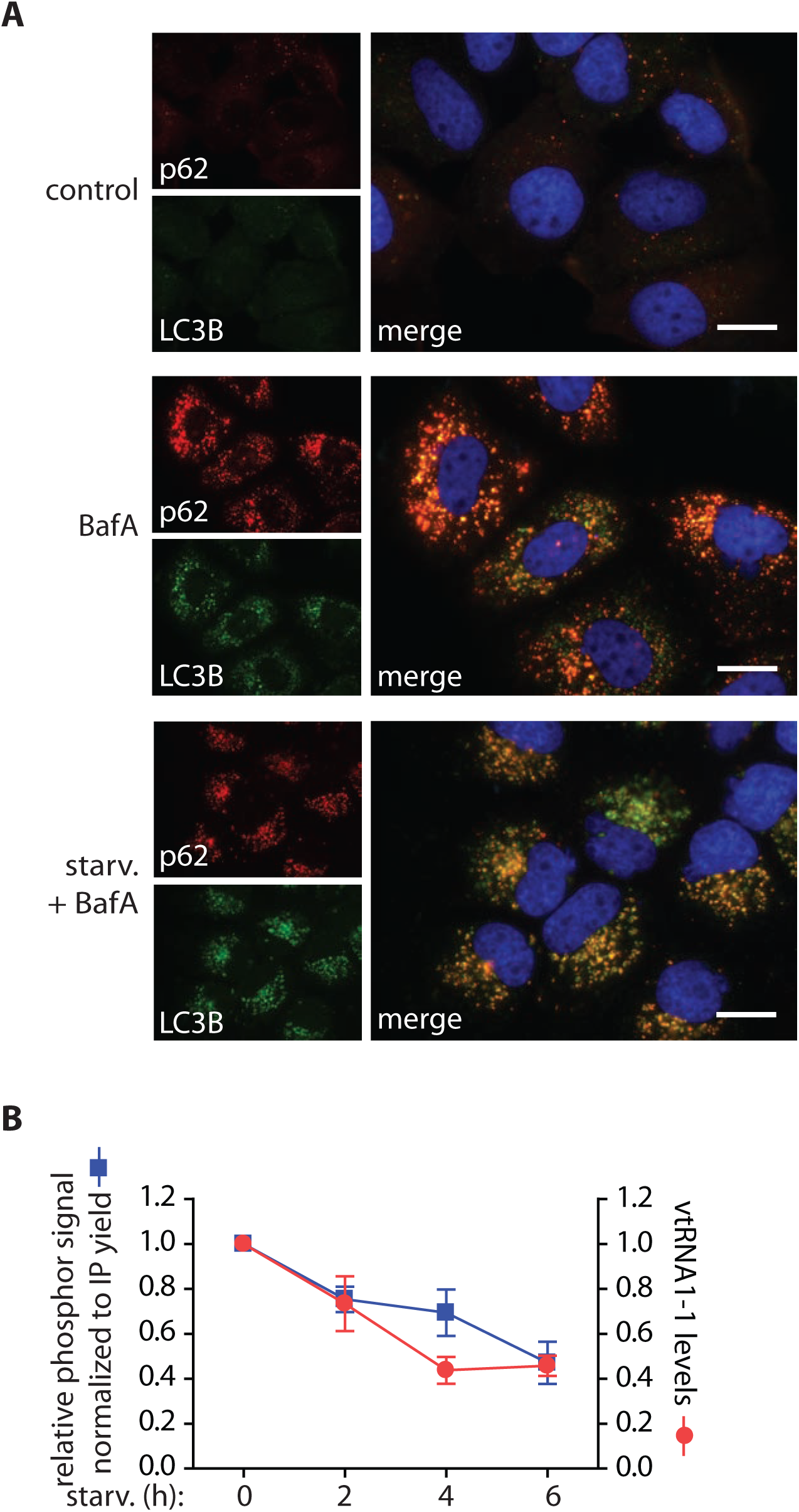
Analysis of p62 localization and RNA binding. (**A**) HuH-7 cells were treated for 4 hours with BafA at 100nM, or starved for 4 hours in the presence of BafA at 100nM. Cells were then fixed with paraformaldehyde and stained with anti-p62 (red), anti-LC3B (green) and DAPI (blue). Representative images are shown; scale bars represent 20 µm. (**B**) Data of quantified RNA-bound p62/total p62 from Fig. 4B is plotted together with the total vtRNA1-1 levels acquired from the HuH-7 cells starved in minimal media for indicated time.

**Table S1.**
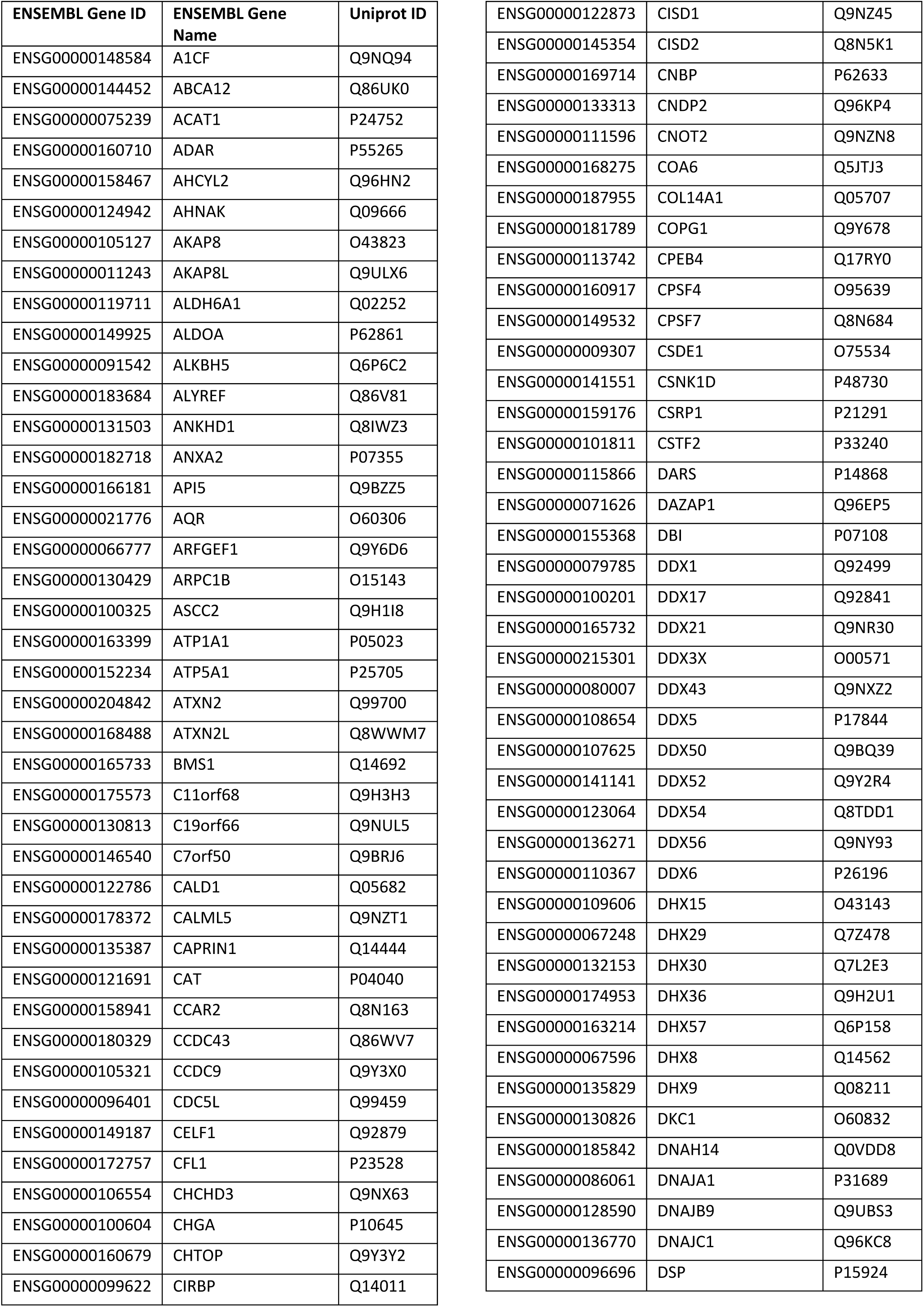
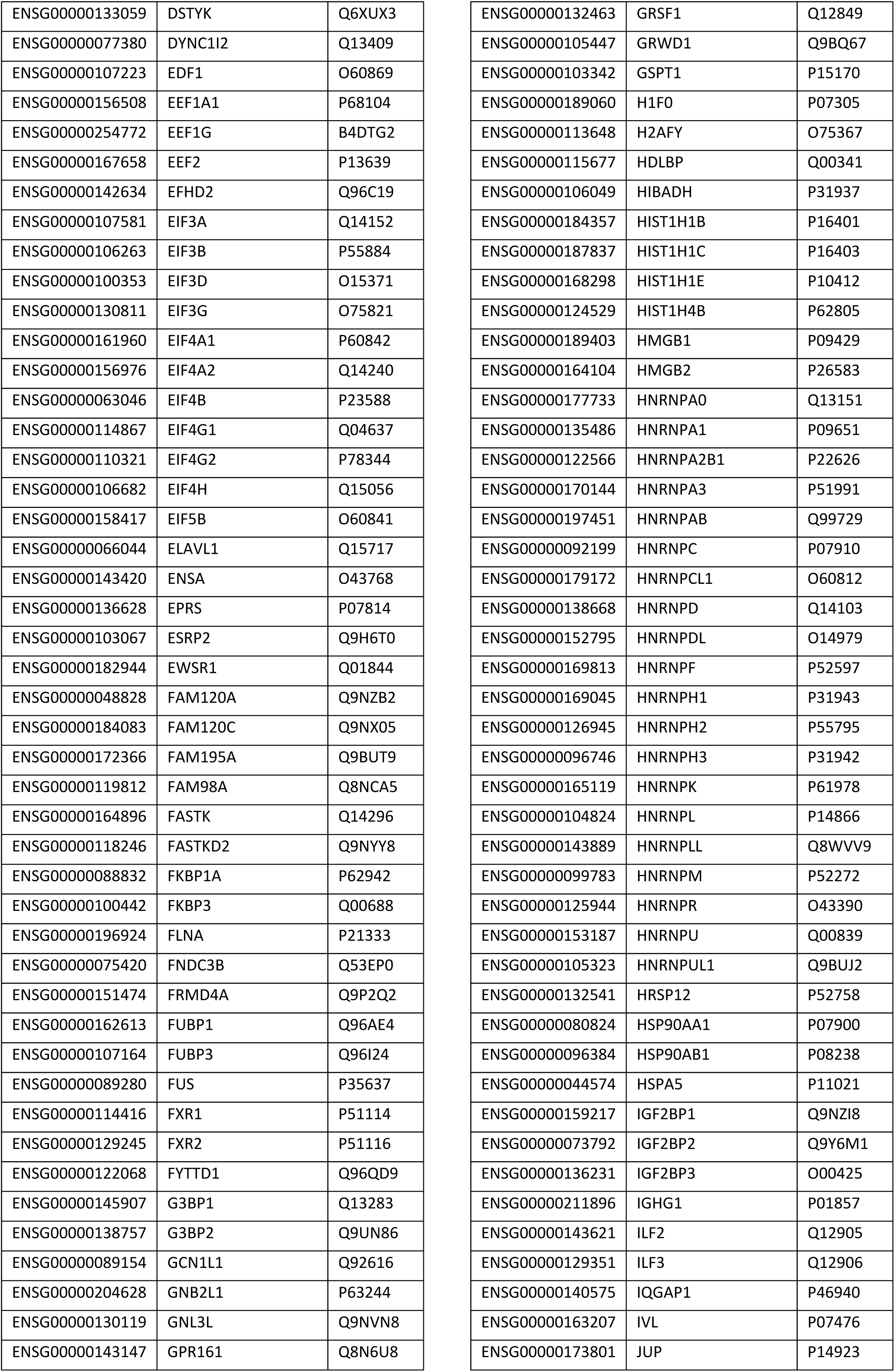
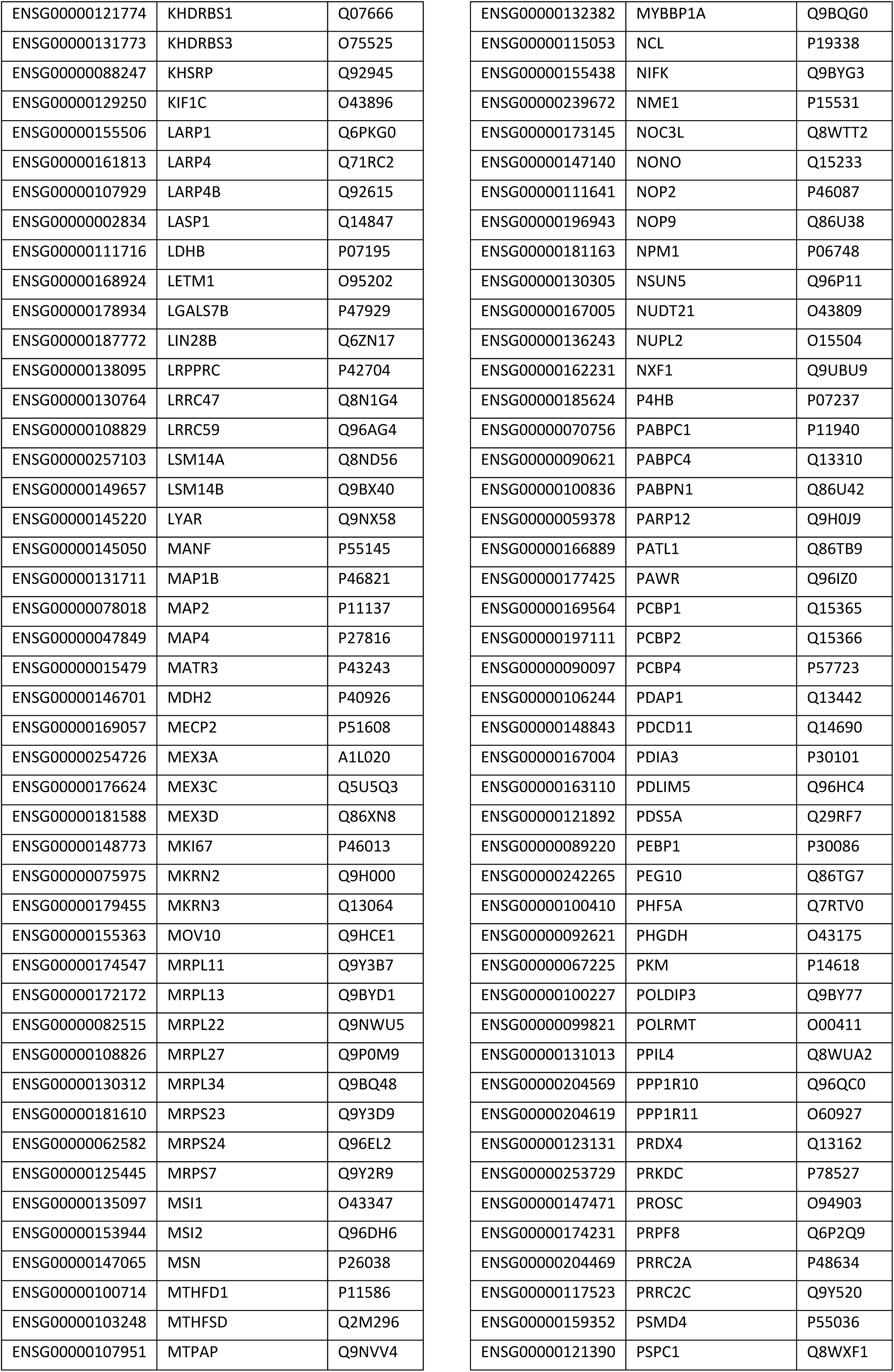
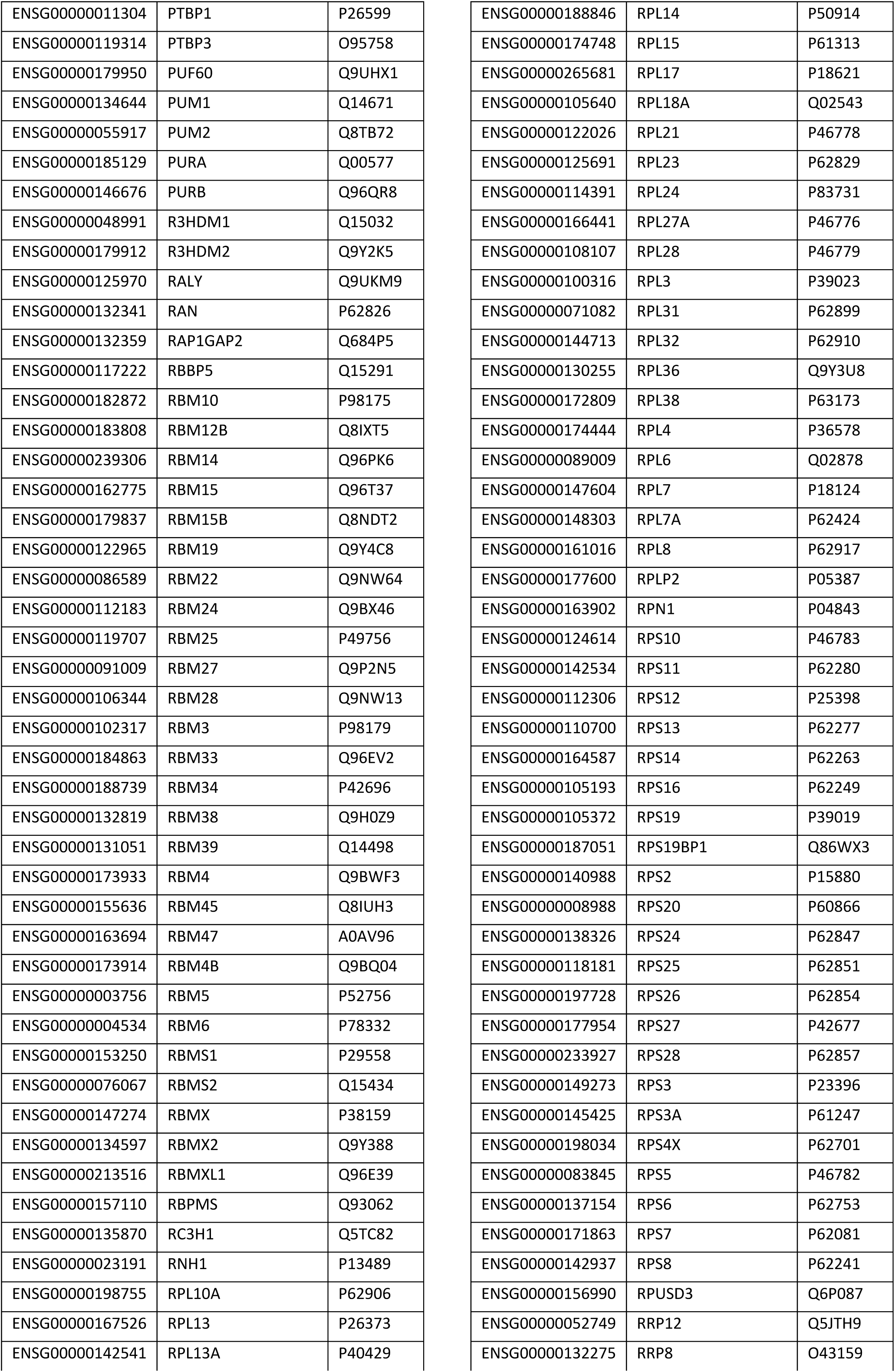
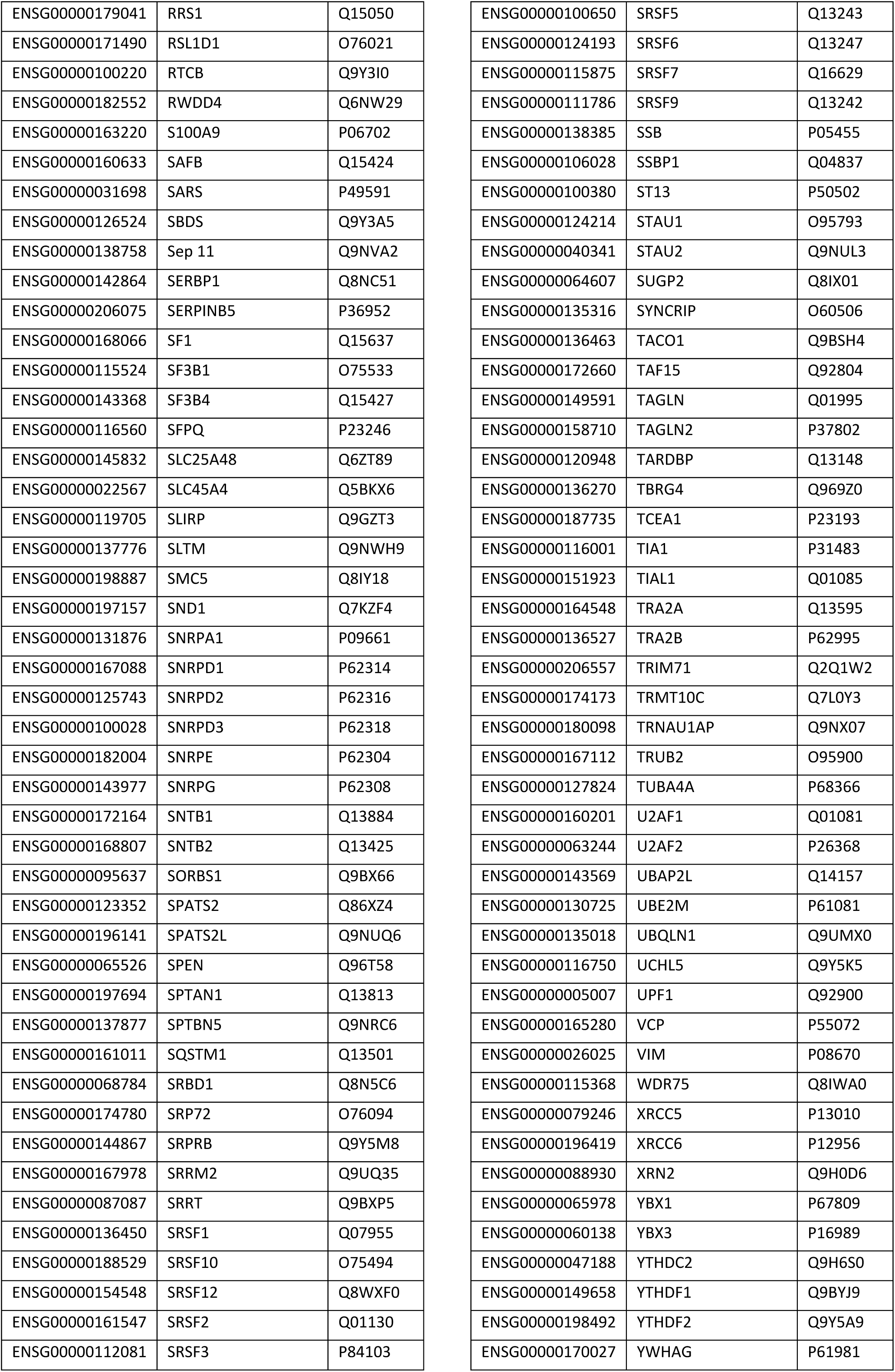
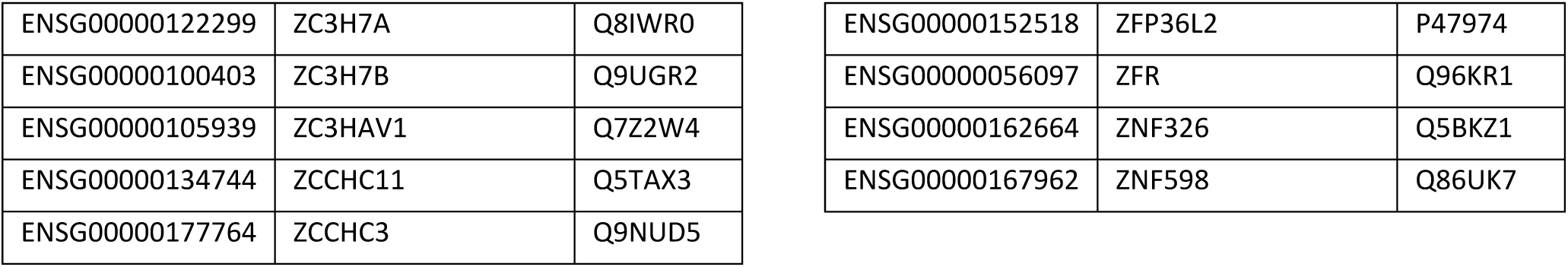
This table lists the proteins found in the HuH-7 RBDmap.

**Table S2.**
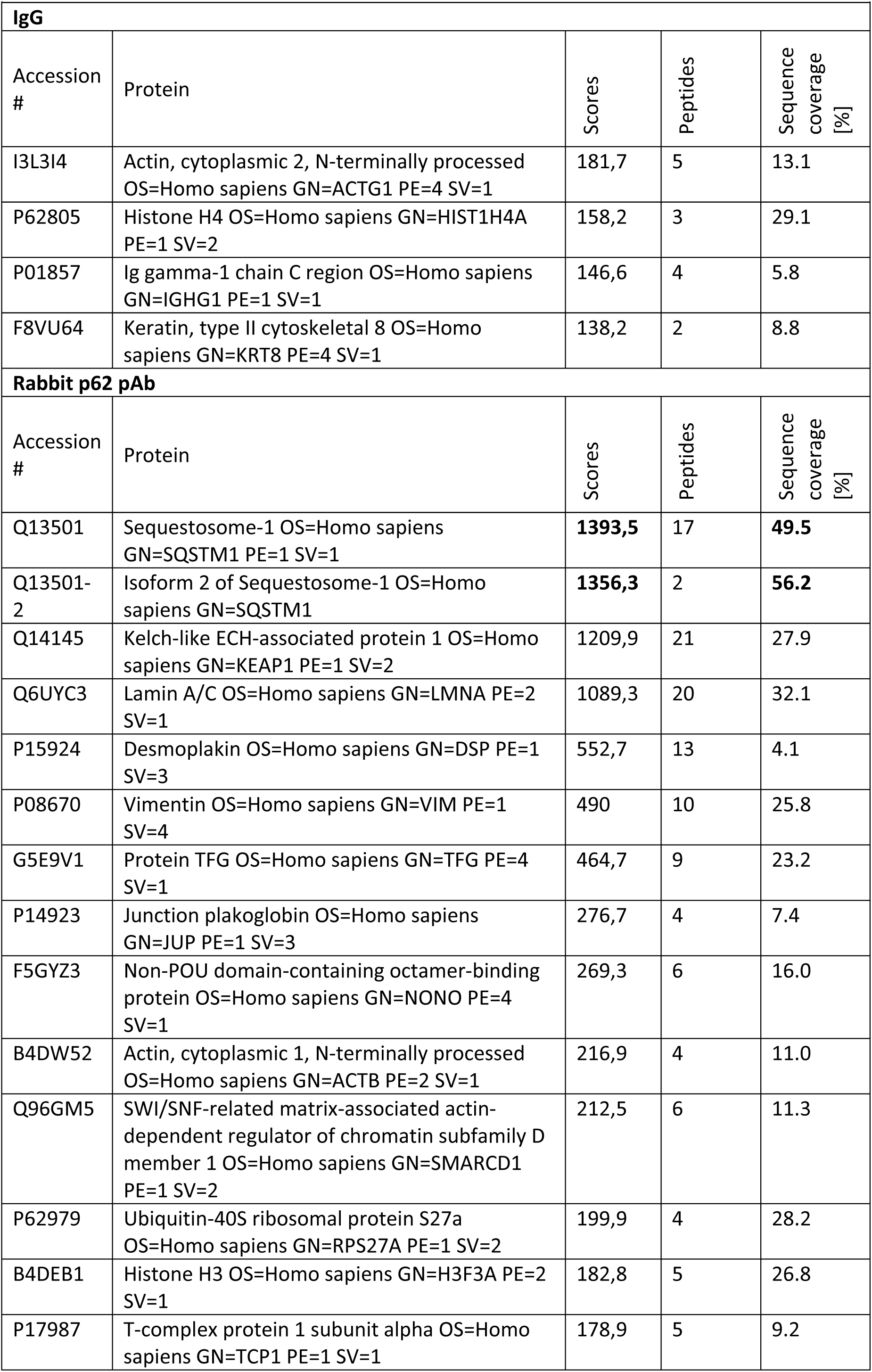
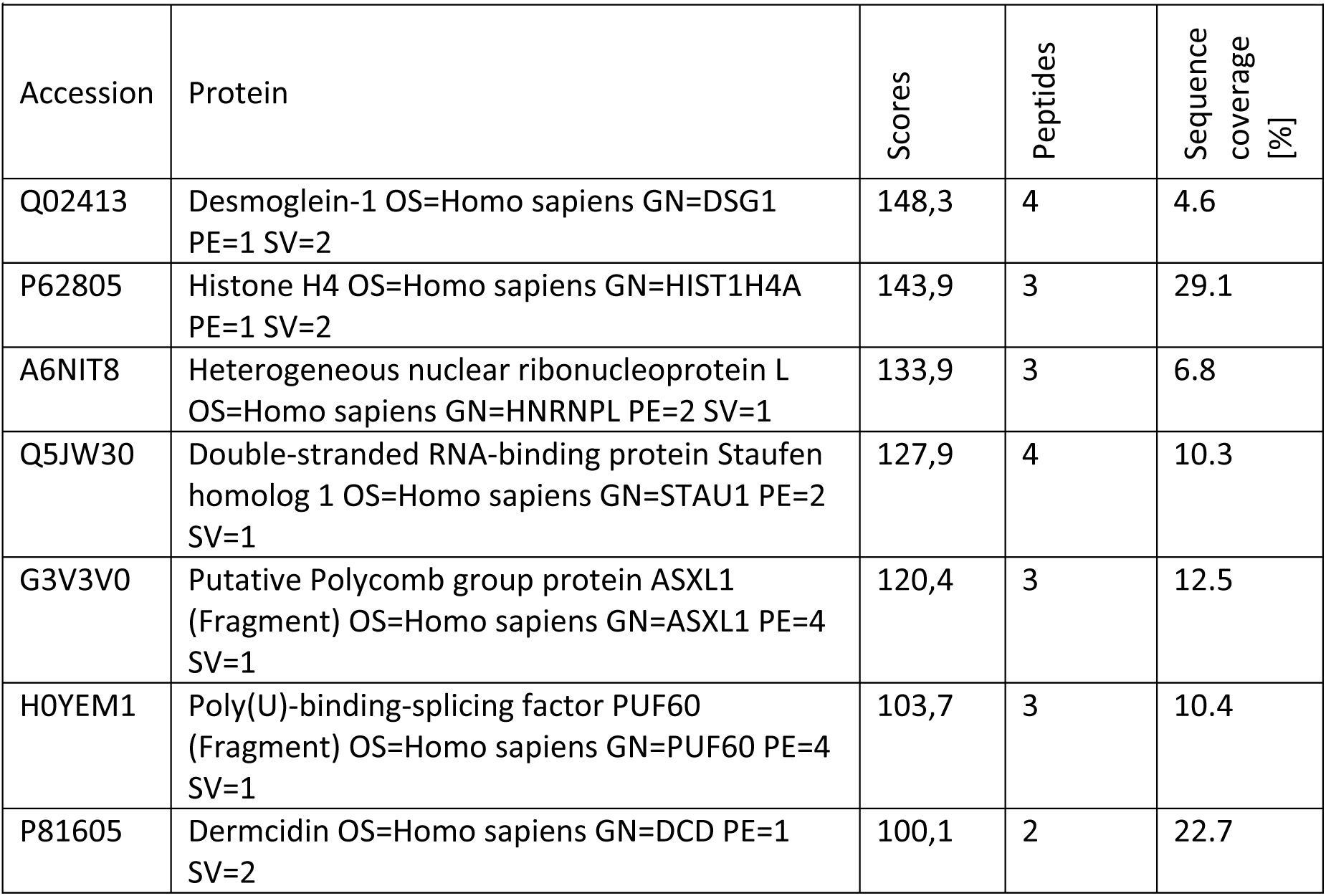
Proteomics analysis of IP eluates after control (IgG) or polyclonal rabbit p62 antibody pulldown excised from SDS-PAGE gel around 60 kDa MW.

**Table S3.** Crosslink sites per RNA type, length and library size normalized. M…Mouse R…Rabbit A / B … Batches pAb – polyclonal antibody, mAb – monoclonal antibody

|  | Total<br>length | Rabbit<br>pAb -B | Rabbit<br>pAb -A | Mouse<br>mAb-A | Mouse<br>mAb -<br>B | Empty<br>beads-A | Empty<br>beads-B | IgG-M-A | IgG-M-B | IgG-R-A | IgG-R-B |
| --- | --- | --- | --- | --- | --- | --- | --- | --- | --- | --- | --- |
| vaultRNA | 377 | 51497,73 | 56495,53 | 45851,25 | 63375,86 | 2954,18 | 10971,78 | 4922,89 | 4814,80 | 1630,78 | 1243,00 |
| tRNAscan | 45214 | 9759,71 | 6495,51 | 9245,27 | 13042,38 | 2863,51 | 4116,78 | 1866,69 | 3378,99 | 1998,86 | 4425,55 |
| Mt_rRNA | 2513 | 3862,84 | 4454,75 | 7803,30 | 4599,45 | 5983,02 | 5102,55 | 7420,48 | 7825,09 | 5137,65 | 5407,76 |
| Mt_tRNA | 1508 | 802,72 | 1197,35 | 3034,74 | 1002,62 | 2954,18 | 3017,24 | 3106,11 | 2206,78 | 3941,06 | 3107,50 |
| rRNA | 59444 | 129,23 | 56,66 | 282,02 | 114,46 | 231,07 | 507,96 | 127,86 | 386,79 | 113,77 | 654,31 |
| misc_RNA | 426141 | 35,02 | 39,41 | 34,78 | 33,43 | 28,53 | 34,94 | 19,49 | 24,85 | 41,84 | 37,39 |
| snRNA | 207473 | 25,47 | 20,96 | 39,09 | 25,70 | 72,92 | 59,81 | 47,28 | 113,74 | 98,78 | 101,64 |
| miRNA | 269361 | 20,22 | 13,60 | 21,39 | 29,76 | 23,09 | 49,14 | 23,62 | 44,93 | 23,59 | 31,31 |
| 3UTR | 27069668 | 2,99 | 4,86 | 3,74 | 3,77 | 2,79 | 2,55 | 3,36 | 2,94 | 2,66 | 1,63 |
| 5UTR | 25804825 | 2,98 | 4,84 | 3,80 | 3,79 | 3,18 | 2,42 | 3,46 | 2,88 | 2,53 | 2,11 |
| CDS | 28815147 | 2,67 | 4,56 | 3,57 | 3,53 | 2,00 | 2,01 | 2,38 | 2,86 | 1,69 | 2,07 |
| protein_coding_exon | 16764974 | 1,35 | 1,60 | 1,25 | 1,21 | 1,33 | 0,94 | 1,58 | 1,21 | 1,22 | 0,67 |
| snoRNA | 163910 | 1,14 | 2,79 | 3,18 | 1,82 | 9,06 | 12,62 | 11,32 | 5,54 | 8,75 | 2,86 |
| transcribed_unitary_pseudogene | 28235 | 0,82 | 0,00 | 1,68 | 0,00 | 0,00 | 0,00 | 0,00 | 0,00 | 0,00 | 0,00 |
| macro_lncRNA | 205012 | 0,79 | 1,44 | 1,97 | 0,68 | 0,91 | 0,00 | 1,29 | 1,48 | 7,00 | 0,00 |
| processed_pseudogene | 8047388 | 0,58 | 1,18 | 1,05 | 1,07 | 1,22 | 1,03 | 0,90 | 1,58 | 0,87 | 1,34 |
| processed_transcript | 25693208 | 0,56 | 0,50 | 0,44 | 0,51 | 0,35 | 0,64 | 0,38 | 0,51 | 0,54 | 0,35 |
| protein_coding_intron | 1229641747 | 0,50 | 0,50 | 0,45 | 0,35 | 0,57 | 0,49 | 0,59 | 0,52 | 0,57 | 0,43 |
| antisense | 116193196 | 0,41 | 0,37 | 0,37 | 0,40 | 0,29 | 0,27 | 0,31 | 0,25 | 0,25 | 0,30 |
| sense_overlapping | 6952261 | 0,28 | 0,27 | 0,28 | 0,15 | 0,20 | 0,54 | 0,32 | 0,48 | 0,44 | 1,21 |
| transcribed_processed_pseudogene | 4749013 | 0,26 | 0,24 | 0,27 | 0,16 | 0,57 | 0,44 | 0,35 | 0,57 | 0,65 | 0,69 |
| 3prime_overlapping_ncrna | 194829 | 0,24 | 0,38 | 0,37 | 0,20 | 0,00 | 2,12 | 1,81 | 0,00 | 0,00 | 0,00 |
| TEC | 1757413 | 0,24 | 0,17 | 0,16 | 0,17 | 0,21 | 0,00 | 0,10 | 0,17 | 0,35 | 0,00 |
| lincRNA | 217616266 | 0,23 | 0,22 | 0,23 | 0,20 | 0,26 | 0,19 | 0,25 | 0,25 | 0,30 | 0,28 |
| transcribed_unprocessed_pseudogene | 15883155 | 0,22 | 0,23 | 0,20 | 0,17 | 0,25 | 0,29 | 0,28 | 0,23 | 0,26 | 0,09 |
| unitary_pseudogene | 4077426 | 0,21 | 0,20 | 0,20 | 0,18 | 0,18 | 0,20 | 0,20 | 0,00 | 0,25 | 0,11 |
| polymorphic_pseudogene | 572666 | 0,08 | 0,06 | 0,00 | 0,00 | 0,32 | 0,00 | 0,46 | 0,00 | 1,79 | 0,00 |
| unprocessed_pseudogene | 12920475 | 0,06 | 0,06 | 0,07 | 0,04 | 0,19 | 0,19 | 0,11 | 0,19 | 0,13 | 0,36 |
| sense_intronic | 4990054 | 0,05 | 0,06 | 0,09 | 0,06 | 0,02 | 0,08 | 0,04 | 0,00 | 0,08 | 0,00 |
| pseudogene | 3547420 | 0,02 | 0,05 | 0,07 | 0,06 | 0,37 | 0,12 | 0,00 | 0,00 | 0,00 | 0,00 |
| IG_C_pseudogene | 9112 | 0,00 | 0,00 | 0,00 | 0,00 | 0,00 | 0,00 | 0,00 | 0,00 | 0,00 | 0,00 |
| IG_D_gene | 851 | 0,00 | 0,00 | 0,00 | 0,00 | 0,00 | 0,00 | 0,00 | 0,00 | 0,00 | 0,00 |
| IG_J_gene | 1198 | 0,00 | 0,00 | 0,00 | 0,00 | 0,00 | 0,00 | 0,00 | 0,00 | 0,00 | 0,00 |
| IG_J_pseudogene | 165 | 0,00 | 0,00 | 0,00 | 0,00 | 0,00 | 0,00 | 0,00 | 0,00 | 0,00 | 0,00 |
| IG_V_pseudogene | 69688 | 0,00 | 0,00 | 0,00 | 0,00 | 0,00 | 0,00 | 0,00 | 0,00 | 0,00 | 0,00 |
| translated_unprocessed_pseudogene | 44659 | 0,00 | 0,00 | 0,00 | 0,00 | 0,00 | 0,00 | 0,00 | 0,00 | 0,00 | 0,00 |
| TR_D_gene | 39 | 0,00 | 0,00 | 0,00 | 0,00 | 0,00 | 0,00 | 0,00 | 0,00 | 0,00 | 0,00 |
| TR_J_gene | 4308 | 0,00 | 0,00 | 0,00 | 0,00 | 0,00 | 0,00 | 0,00 | 0,00 | 0,00 | 0,00 |
| TR_J_pseudogene | 299 | 0,00 | 0,00 | 0,00 | 0,00 | 0,00 | 0,00 | 0,00 | 0,00 | 0,00 | 0,00 |
| TR_V_gene | 61855 | 0,00 | 0,00 | 0,00 | 0,00 | 0,00 | 0,00 | 0,00 | 0,00 | 0,00 | 0,00 |
| TR_V_pseudogene | 13064 | 0,00 | 0,00 | 0,00 | 0,00 | 0,00 | 0,00 | 0,00 | 0,00 | 0,00 | 0,00 |
| vaultRNA_pseudogene | 102 | 0,00 | 0,00 | 0,00 | 0,00 | 0,00 | 0,00 | 0,00 | 0,00 | 0,00 | 0,00 |
| TR_C_gene | 638633 | 0,00 | 0,00 | 0,04 | 0,00 | 0,00 | 0,00 | 0,00 | 0,00 | 0,00 | 0,00 |
| IG_C_gene | 1898345 | 0,00 | 0,00 | 0,00 | 0,01 | 0,00 | 0,00 | 0,00 | 0,00 | 0,00 | 0,00 |
| IG_V_gene | 441901 | 0,00 | 0,00 | 0,05 | 0,05 | 0,00 | 0,00 | 0,00 | 0,00 | 0,00 | 0,00 |

